# Salicylic acid-induced alkalinization of the apoplast requires *TRANSMEMBRANE KINASE 1* and results in growth attenuation

**DOI:** 10.64898/2025.12.02.691772

**Authors:** Jonas Müller, Kaltra Xhelilaj, Marjorie Guichard, Sabrina Kaiser, Guido Grossmann, Raimund Tenhaken, Julien Gronnier, David Scheuring

**Author notes:** Correspondence, **Corresponding author** Correspondence to David Scheuring. The author responsible for distribution of materials integral to the findings presented in this article is David Scheuring.

## Abstract

The phytohormone salicylic acid (SA) has a key role in regulating plant growth and stress response. In the past, most of the growth-related SA functions have been explained by crosstalk with the master growth regulator auxin. By affecting polarity of auxin transporters, SA changes auxin distribution in the root and inhibits growth of the main root and activates lateral root formation. However, only recently there is evidence emerging that SA impacts growth processes independently of nuclear auxin signalling, possessing yet unknown mechanistic functions. Here we show that SA activity depends on *TRANSMEMBRANE KINASE 1* (*TMK1*) resulting in apoplast alkalinization and growth restriction. SA treatment prevents phosphorylation and activation of plasma membrane (PM) H^+^-ATPases at the cell surface and does not depend on the auxin receptor Auxin Binding Protein 1 (ABP1). We suggest that alkalinization of the apoplast by SA serves as mechanism to balance stress response and growth.

## Introduction

Salicylic acid (SA) is a key phytohormone regulating plant immunity and stress responses, but it also modulates plant growth and development. SA’s effects on growth are dose-, duration-, and tissue-dependent: low SA levels promote growth, while elevated endogenous SA correlates with growth inhibition (Friedrich *et al*., 1995; Bowling *et al*., 1997; Rate *et al*., 1999; Wildermuth *et al*., 2001). However, the mechanisms underlying SA-induced growth restriction remain unclear. Although the immune regulator Nonexpresser of PR genes 1 (NPR1) is essential for SA-mediated defense, it is not required for growth inhibition (Tan *et al*., 2020; Müller *et al*., 2025). However, SA can directly bind to protein phosphatase 2A (PP2A) subunits, affecting the auxin transporter PIN2 by promoting its hyperphosphorylation which in turn abolishes polarity of the transporter, inhibiting root growth and gravitropic response eventually (Tan *et al*., 2020).

SA – auxin crosstalk as basis of SA function has been proposed earlier (Pasternak *et al*., 2019) but there is also evidence for auxin-independency. Firstly, it is known that PP2A acts on different substrates and *pp2aa1* mutation led to increased SA sensitivity and not resistance (Tan *et al*., 2020), questioning this mechanism to be the basis for root growth inhibition. Secondly, it has been demonstrated that SA inhibits V-ATPase activity independently of canonical nuclear (TIR/AFB) auxin signalling, leading to increased vacuolar pH (Müller *et al*., 2025). In addition, the accompanying changes of vacuolar morphology and SNARE abundance are opposite of the effect induced by auxin: While auxin induces constricted and smaller vacuoles and accumulation of tonoplast soluble N-ethylmaleimide-sensitive-factor attachment receptor (SNAREs) (Löfke *et al*., 2015; Scheuring *et al*., 2016), SA application leads to homotypic vacuole fusion with largely unchanged size along with a decrease in the abundance of the tonoplast SNARE SYP21 (Müller *et al*., 2025). Thus, an auxin independent mechanism has been proposed to explain SA-induced growth restriction.

In general, the pH of plant cells is strictly controlled in each compartment. The pH in the apoplast has been directly related to cell wall stiffness which is crucial for cell elongation and eventually growth. The so-called “acid growth theory” includes the activation of plasma membrane (PM) H^+^-ATPases by auxin and subsequent acidification of the apoplast (Du *et al*., 2020). By this, pH-responsive non-enzymatic proteins from the expansin family are activated (McQueen-Mason *et al*., 1992), mediating cell wall loosening. Cell elongation is subsequently achieved by communication with and swelling of the vacuole as well as deposition of new cell wall material (Majda & Robert, 2018; Kaiser & Scheuring, 2020). By targeting PM H^+^-ATPases, several phytohormones inhibit apoplast acidification as a mean to control cell elongation and eventually regulate plant growth (Gámez-Arjona *et al*., 2022).

Here, we show that SA treatment leads to changes of cytosolic and apoplastic pH independent of canonical nuclear auxin signalling. In dependence of *TRANSMEMBRANE KINASE 1* (*TMK1*) at the PM, SA inhibits phosphorylation and hence activation of PM H^+^-ATPases, resulting in a decreased H^+^ export and alkalinization of the apoplast. Notably, this effect is independent of the SA receptors NPRs and PP2A, suggesting a yet undescribed mechanism to inhibit growth. Our data indicates that SA-induced alkalinization of the apoplast by inhibition of PM H^+^-ATPases activity blocks cell wall loosening required for cell elongation. The independence of this mechanism from NPR-based immunity suggests an additional SA function, potentially integrating stress responses into plant growth and development.

## Results

Previously, we demonstrated that SA inhibits V-ATPase activity causing a significant increase of vacuolar pH (Müller *et al*., 2025). This suggests an intracellular shift in proton concentration, potentially accumulating in the cytosol. To follow this, we employed circular permutated YFP (cpYFP) as cytosolic pH sensor (Schwarzländer *et al*., 2011) in epidermal cells from the transition zone of Arabidopsis roots (Fig. 1A). Ratiometric measurements revealed a rapid but lasting decrease of cytosolic pH already after 30 min of SA treatment (Fig. 1B).

**Figure 1.**
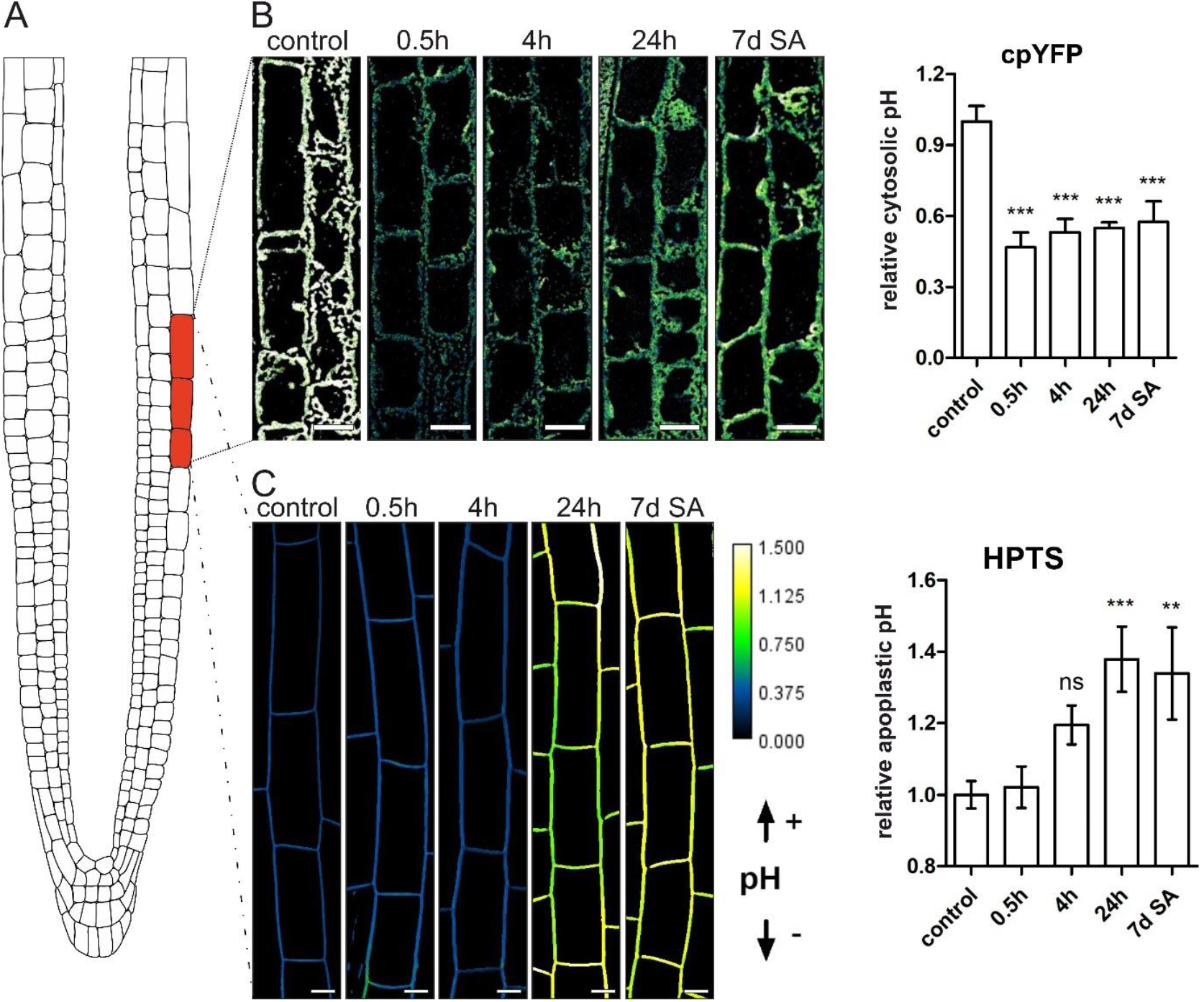
SA reduces cytosolic pH while apoplastic pH is increased. **A)** Region of the Arabidopsis root used for microscopic analysis. **B)** Seedlings of the cytosolic pH marker line cpYFP were grown on ½MS plates and treated with 50 µM SA for 0.5h, 4h, 24h, or 7 days. Ratiometric images were generated by dividing the λ_ex_488 fluorescence intensity by the λ_ex_405 intensity and subsequently quantified. SA-treated samples were normalized to the corresponding control (n control = 29, n 0.5h = 16, n 4h = 18, n 24h = 16, n 7d SA = 13). Scale bar: 15 µm. Data was analyzed using ANOVA with Tukey post-hoc test. \*\*\**P* ≤ 0.001. **C)** Col-0 seedlings were grown on ½MS plates and treated with 50 µM SA for 0.5h, 4h, 24h, and 7 days and stained with HPTS. Ratiometric images and quantification were done by dividing the λex 458 intensity by the λex 405 intensity (n control = 35 roots, n 0.5h = 21, n 4h = 15, n 24h = 23, n 7d SA = 13). Data was analyzed using ANOVA with Tukey post-hoc test. \*\**P* ≤ 0.01, \*\*\**P* ≤ 0.001. Scale bar: 4 µm.

Since accumulation of protons in the cytosol could result from inhibition of proton efflux as well, we measured the pH in the apoplast, using the dye 8-hydroxypyrene-1,3,6-trisulfonic acid trisodium salt (HPTS) (Barbez *et al*., 2017). Analyzing the same region of the root, we found an SA-mediated increase in apoplastic pH which was highly significant after 24h (Fig. 1C). Acidification of the apoplast is an active process, executed by PM-residing H^+^-ATPases (Arabidopsis H^+^-ATPases, AHAs) (Falhof *et al*., 2016). To test if SA interferes with PM H^+^-ATPase activity, we used SA in parallel with the fungal toxin fusicoccin, a potent activator of PM H^+^-ATPase (Baunsgaard *et al*., 1998). Treatment with fusicoccin alone decreased apoplastic pH as expected, while prolonged SA treatment resulted in a significantly increased pH (Fig. 2A). Co-treatment with fusicoccin abolished the SA-induced pH increase, suggesting a SA-induced inhibition of PM H^+^-ATPase activity.

**Figure 2.**
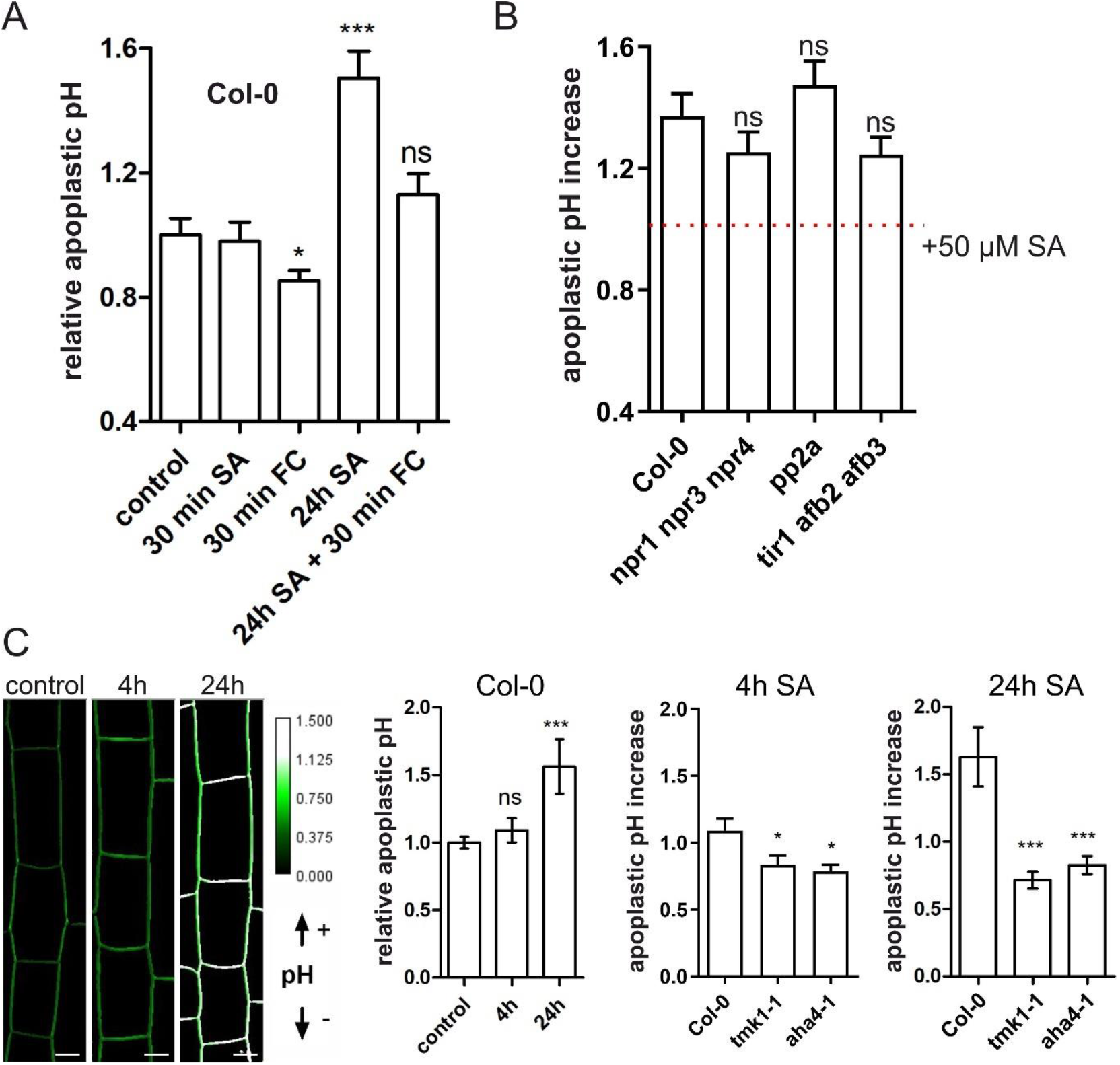
TMK1 and AHA4 mutants are resistant to SA-mediated increase of apoplastic pH. **A)** Col-0 seedlings were grown on ½MS plates for 7 days and subsequently treated with SA, fusicoccin (FC), or a combination of both. Relative apoplastic pH was quantified using HPTS staining by normalizing the λe_x_458/λ_ex_405 ratio to the DMSO control. (n control = 13, n 30 min SA = 5, n 30 min FC = 15, n 24h SA = 4, n 24h SA + 30 min FC = 14). Data was analyzed using one-way ANOVA with Tukey post-hoc test. \**P* ≤ 0.05, \*\*\**P* ≤ 0.001. **B)** Apoplastic pH increase was measured by staining 7-day-old Col-0 and mutant seedlings with HPTS and quantifying the λ_ex_458/λ_ex_405 ratio. Additionally, seedlings were grown for 24 hours on SA-supplemented medium, and the λ_ex_458/λ_ex_405 ratio was normalized to the corresponding untreated control. (n Col-0 = 35, n Col-0 SA = 27, n *npr1 npr3 npr4* = 13, n *npr1 npr3 npr4* SA = 13, n *pp2a* = 24, n *pp2a* SA = 26, n *tir1 afb2 afb3* = 26, n *tir1 afb2 afb3* SA = 29). The dotted line displays normalization. Data were analyzed with Student’s t-test. **C)** Col-0 seedlings were grown on ½ MS plates for 6 days and then either treated with SA for 4 hours or transferred to ½MS+ plates supplemented with SA for 24 hours. Relative apoplastic pH was quantified using HPTS staining by normalizing the λ_ex_458/λ_ex_405 fluorescence ratio to the DMSO control. (n control = 25, n 4h = 6, n 24h = 13). Data was analyzed using one-way ANOVA with Tukey post-hoc test. \*\*\**P* ≤ 0.001. **D)** 7-day old Col-0, *tmk1-1* and *aha4-1* seedlings were stained with HPTS and the λ_ex_458/λ_ex_405 ratio quantified. For normalization the λex λ_ex_458/λ_ex_405 ratio of Col-0, *tmk1-1* and *aha4-1* seedlings treated for 4 hours with SA (n control = 7, n control SA = 6, n *tmk1-1* = 6, n *tmk1-1* SA = 8, n *aha4-1* = 6, n *aha4-1* SA = 6) and 24 hours with SA (n control = 20, n control SA = 18, n *tmk1-1* = 20, n *tmk1-1* SA = 20, n *aha4-1* = 22, n *aha4-1* SA = 24) were quantified. Data was analyzed with Student’s t-test. \**P* ≤ 0.05, \*\*\**P* ≤ 0.001.

To investigate whether the SA effect on apoplastic pH depends on canonical NPR-dependent signalling or the reported auxin crosstalk (Pasternak *et al*., 2019; Tan *et al*., 2020), we measured pH in the SA-receptor triple mutant *npr1 npr 3 npr4*, a mutant of the reported SA-binding phosphatase PP2A and the auxin receptor triple mutant *tir1 afb2 afb3*. None of the lines displayed any resistance against SA-induced increase of apoplastic pH (Fig. 2B). Since it has been shown that TRANSMEMBRANE KINASE 1 (TMK1) activates PM H^+^-ATPases by phosphorylation which leads to apoplast acidification (Li *et al*., 2021; Lin *et al*., 2021), we tested *tmk1-1* and *aha4-1*, which is abundant in the root epidermis (Vitart *et al*., 2001) for pH changes in the apoplast. After 24h SA treatment apoplastic pH increased in the Col-0 wildtype whereas *tmk1-1* and *aha4-1* both showed resistance (Fig. 2D). To test if the observed resistance translates into better growth, we used a vertical set-up to measure root length. To this end we used SPIRO, the Smart Plate Imaging Robot (Ohlsson *et al*., 2024) and transferred 6 day old seedlings onto SA and control plates. Using the SPIRO, we hourly captured growth and calculated the additional growth in percent, 5h after transfer (Fig. 3A). FC treatment led to significantly better growth, while SA reduced growth (Fig. 3B). Although the additional growth mean was closer to the control upon FC and SA co-treatments, the SA-induced growth reduction could not be rescued.

**Figure 3.**
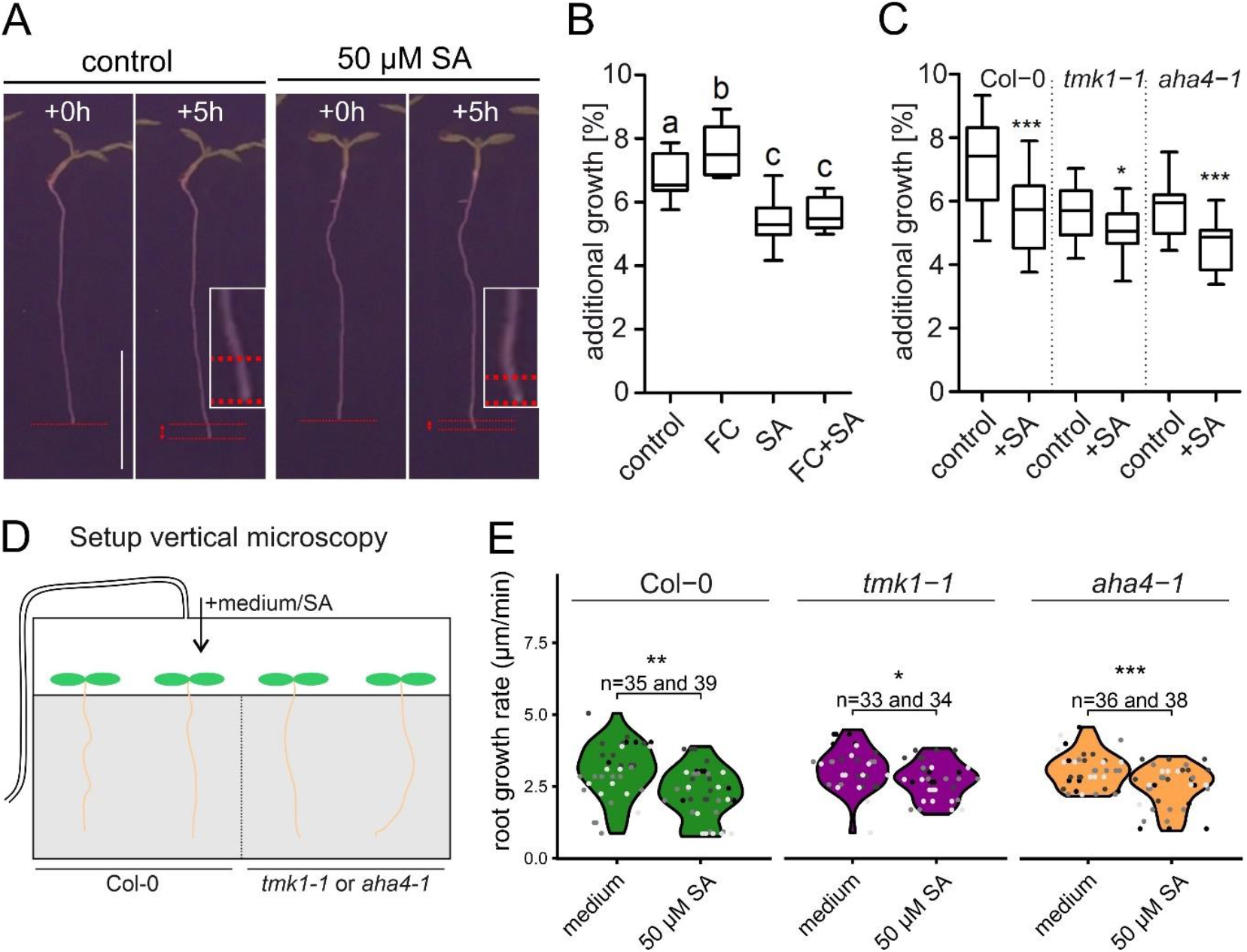
TMK1 is partially resistant to SA-mediated root growth inhibition. **A)** Representative images of how additional growth was evaluated. Col-0 seedlings were grown for 6 days on ½MS plates. After 5 hours the additional growth was measured (red arrow between red dotted lines) and taken relative to the whole root length. Scalebar=1 cm. **B)** Six-day-old Col-0 seedlings were grown on ½MS plates and then transferred to control medium, 50 µM SA, 5 µM fusicoccin (FC), or a combination of both. Additional growth was subsequently quantified (n control = 12, n FC = 13, n SA = 15, n FC+SA = 12). Change in letter equals p≤0.001. **C)** Six-day-old seedlings of Col-0, *tmk1-1*, and *aha4-1* were grown on ½MS plates and then either left on control plates or transferred to ½MS plates supplemented with 50 µM SA. Root growth was measured after 5 hours and expressed relative to the total root length (n Col-0 = 48, n Col-0 SA = 49, n *tmk1-1* = 34, n *tmk1-1* SA = 37, n *aha4-1* = 36, n aha4-1 SA = 32). Data is presented in whisker plots with statistical analysis performed using Student’s t-test. *p≤0.05, ***p≤0.001. **D)** Schematic illustration of the RoPod setup used for root growth rate measurements. Seedlings were grown in microscope-compatible RoPod chambers for 6 days. Subsequently, ½ MS medium supplemented with 50 µM SA was injected into one RoPod, while the control RoPod received ½ MS medium containing ethanol. Root growth was recorded over the following 5 hours. **E)** Col-0, *tmk1-1, and aha4-1* seedlings were grown for 6 days in the RoPod system, and root growth rates were monitored using time-lapse microscopy. Liquid ½MS medium as control or ½ MS medium supplemented with 50 µM SA was injected into the RoPods, and root growth rates were measured for the following 5 hours. Data are presented in violin plots with statistical analysis performed using Student’s t-test, with a Welch approximation for unequal variance. \**P* ≤ 0.05, \*\*\**P* ≤ 0.001. The root population size for each group is indicated on top of the individual violin plots. Each dot color represents an independent experimental run.

However, as we analyzed growth upon SA treatment (additional growth) using *tmk1-1* and *aha4-1*, we found *tmk1*-1 to be partially resistant against the SA effect (Fig. 3C and Fig. S1). For confirmation, we used the RoPod system (Guichard *et al*., 2024) for time-lapse imaging of root growth on a vertical mount microscope. Different Arabidopsis genotypes were grown for 5-6 days in the RoPod chambers and subsequently treated at the microscope. Equal volumes of control medium and SA-containing medium were injected and the root growth rate measured (Fig. 3D). In comparison to the Col-0 control, the root growth rate of *tmk1-1*, but not *aha4-1*, was significantly less inhibited by SA (Fig. 3E).

So far, our data indicated that SA interferes with PM H^+^-ATPase activation by TMK1. To assess this directly, we measured PM H^+^-ATPase activity on isolated microsomal fractions from seedlings treated for 5h with SA (50 µM) and 1h with FC (1 µM FC). In comparison to the mock-treated seedlings, SA treatment significantly reduced PM H^+^-ATPase activity while the positive control (FC treatment), as expected, showed higher activity (Fig. 4A). To investigate PM H^+^-ATPase activation, the phosphorylation status of AHA was analyzed. To this end, an antibody raised against phosphorylated threonine at the AHA residue 947 (pT947) was employed (Lin *et al*., 2021). Phosphorylation status of PM H^+^-ATPase was then quantified from the ratio pT947/AHA and is expressed relative to the phosphorylation level of the mock control (Fig. 4B). In line with inhibition of PM H^+^-ATPase activity, SA reduced phosphorylation of AHA by more than half (0.4) and FC treatment as positive control increased phosphorylation 3.2-fold.

**Figure 4.**
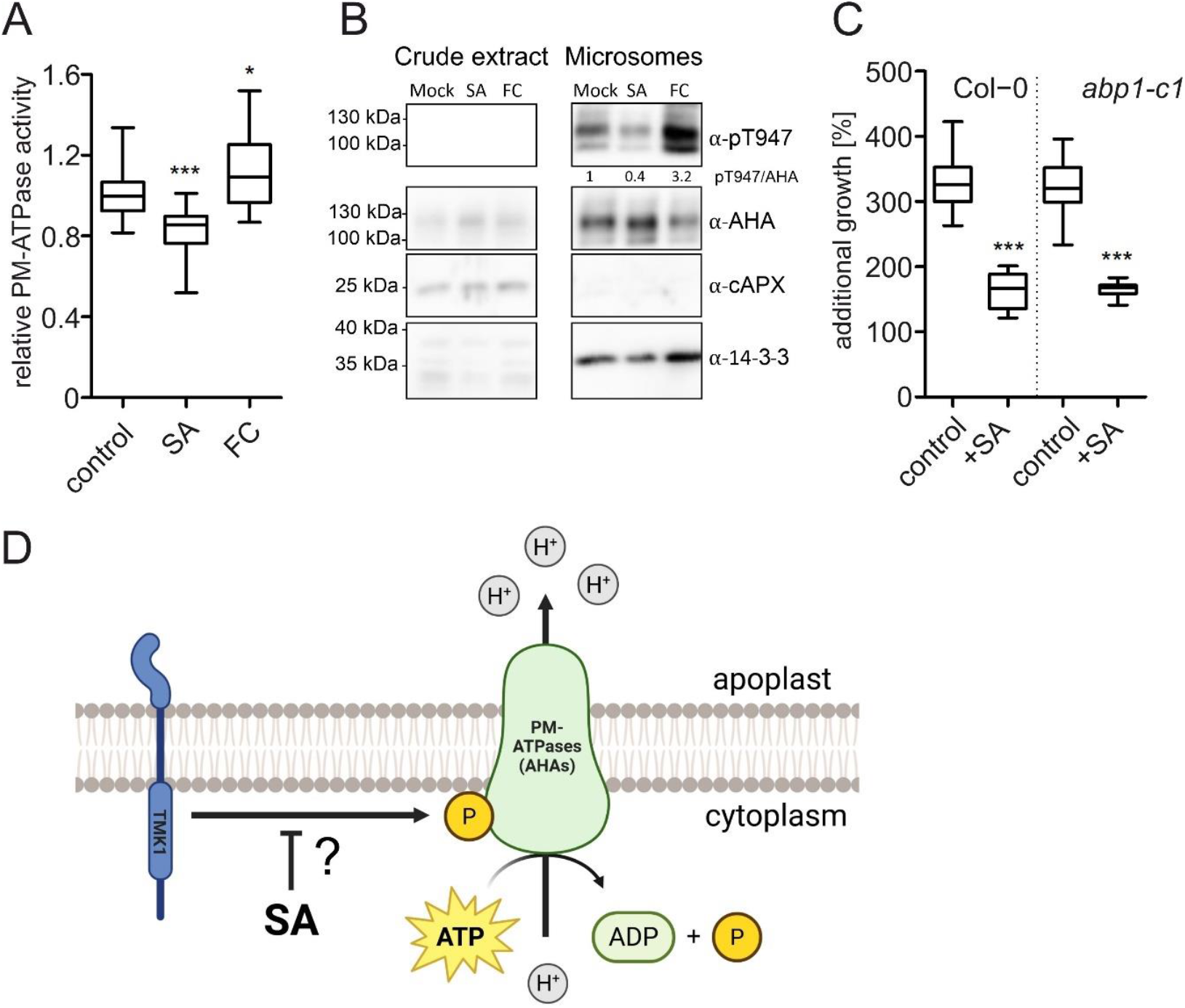
SA reduces PM H^+^-ATPase activity. **A)** Col-0 seedlings were grown on solid ½ MS medium for 4 days and subsequently transferred to liquid ½ MS medium for two weeks. Seedlings were treated with 50 µM SA for 5 hours, with 1 µM fusicoccin (FC) for 1h serving as a positive control, or with DMSO as a mock treatment. Microsomal fractions of roots were isolated, and the ATP hydrolytic activity of PM vanadate-sensitive H^+^-ATPases was determined by quantifying phosphate–malachite complexes as a proxy for released inorganic phosphate (n control = 18, n SA = 18, n FC = 18). Three biological replicates have been conducted. Data is presented in whisker plots with statistical analysis performed using Student’s t-test. *p≤0.05, ***p≤0.001. **B)** For Western blot analysis of AHA phosphorylation, an antibody recognizing the phosphorylated T947 residue of AHA was used and compared with an anti-AHA antibody. The phosphorylation status of the plasma membrane H^+^-ATPase was quantified as the pT947/AHA ratio and expressed relative to the phosphorylation level of the mock control. For additional controls, antibodies against cAPX (marker for purification success) and 14-3-3 were used, both of which are known regulators of plasma membrane H^+^-ATPase activity. **C)** Col-0 and *abp1* seedlings were grown on ½MS plates for six days. For additional growth quantification, seedlings were then transferred for an additional 5 days either to ½MS plates as control or plates supplemented with 50 µM SA (n Col-0 control = 10, n Col-0 SA = 11, n *abp1* control = 10, n *abp1* SA = 11). Data is presented in whisker plots with statistical analysis performed using Student’s t-test. ***p≤0.001. **D)** Working hypothesis: SA-induced apoplast alkalinization is mediated by inhibition of AHA phosphorylation. This suppression could be potentially achieved by blocking TMK1 function by SA.

Since TMK1-based activation of AHAs is central to the rapid auxin signalling pathway, we tested involvement of AUXIN BINDING PROTEIN 1 (ABP1), which is known to interact with TMK1 (Li *et al*., 2021; Lin *et al*., 2021; Friml *et al*., 2022; Fiedler & Friml, 2023). Again, we used the SPIRO to compare additional root growth upon SA treatment of *abp1-c1* and the Col-0 control (Fig. 4C). In contrast to the partial resistance *tmk1-1* showed against SA (Fig. 3C), *abp1-c1* was indistinguishable from the control (Fig. S2). This suggests that the SA effect on the TMK1-AHA module functions independently from the auxin receptor ABP1.

## Discussion

In contrast to its role in stress response, the role of SA in regulating growth and development just recently came into focus. Interestingly, inhibition of root growth by SA has been demonstrated to be independent of NPRs (Tan *et al*., 2020; Müller *et al*., 2025) and the inhibitory effect of SA has been explained by auxin crosstalk or even purely antagonistic effects (Rawat & Laxmi, 2025; Tian *et al*., 2025). However, evidence emerged that some of the SA functions in respect to growth are independent of nuclear auxin signalling and additional signalling pathways have been proposed (Müller and Scheuring, 2025). One of the functions independent from canonical TIR/AFB signalling described is the inhibition of V-ATPase activity resulting in increased vacuolar pH which is detrimental for plant growth (Müller *et al*., 2025). This SA function does not rely on nuclear auxin signalling as the *tir1 afb2 afb3* mutant behaved wildtype-like and in addition no elevated auxin levels in the root could be detected upon SA treatment.

Notably, also the increase of apoplastic pH induced by SA presented here is independent of nuclear auxin signalling (Fig. 2). Auxin has been reported to have antagonistic effects on apoplastic pH depending on whether nuclear or cell surface signalling is triggered. While intracellular (nuclear) auxin signalling leads to alkalinization of the apoplast (Barbez *et al*., 2017; Li *et al*., 2021), cell surface signalling involving TRANSMEMBRANE KINASE 1 (TMK1) results in apoplast acidification (Li *et al*., 2021; Lin *et al*., 2021). Although increase in apoplastic pH and growth inhibition in roots is dependent on the intracellular TIR1/AFB auxin receptors, the effect is non-transcriptional and not yet fully understood (Fendrych *et al*., 2018; Li *et al*., 2021). However, since the observed SA-induced alkalinization of the apoplast is independent of TIR/AFB (Fig. 2), we hypothesized that cell surface-based auxin signalling might be involved. This often called “rapid auxin signalling” relies on the auxin receptor ABP1 at the cell surface and its co-receptor TMK1 (Friml *et al*., 2022; Fiedler & Friml, 2023). The ABP1-TMK1 module has been demonstrated to mediate activation of PM H^+^-ATPases (AHAs) by phosphorylation (Li *et al*., 2021; Lin *et al*., 2021). In roots, this antagonizes the growth-inhibiting TIR/AFB-dependent apoplast alkalinization, allowing dynamic growth regulation (Li *et al*., 2021). Considering that SA prevents AHA phosphorylation, the TMK1-AHA module is inhibited thereby preventing acidification of the apoplast. This translates into significantly reduced root growth inhibition of *tmk1-1* upon SA application while ABP1 does not seem to participate (Fig. 3 and 4). In line with this, ABP1-independent functions of TMKs have been proposed: vasculature formation-related phenotypes of single *tmk* mutants and *abp1* were comparable, but higher-order *tmk* mutants display strong additional phenotypes (Friml *et al*., 2022).

Just recently, it has been demonstrated that the ABP1-TMK1 module (together with ABP-like 3) is responsible for the phosphorylation of the auxin efflux transporter PINFORMED 2 (Rodriguez *et al*., 2025). Following gravistimulation, auxin induces autophosphorylation of TMK1 which leads to interaction with PIN2 resulting in phosphorylation and stabilization of asymmetric PIN2 distribution. Given that SA reduces AHA phosphorylation, it seems possible that SA treatment also interferes with PIN2 phosphorylation and hence, polarity. Indeed, loss of PIN2 polarity upon SA treatment has been reported (Tan *et al*., 2020; Müller *et al*., 2025). However, this was attributed to direct SA binding to and inhibition of PP2A, resulting in hyperphosphorylation of PIN2 and reduced growth (Tan *et al*., 2020). Notably, the *pp2a* mutant was not resistant in response to SA but hypersensitive. Therefore, it seems possible that by targeting AHA activity (this study) and PP2A (Tan *et al*., 2020), SA impacts PIN2 localization twice which might explain the observed hypersensitivity.

Independent of how apoplastic pH increases, the effect alone is detrimental for plant growth according to the *acid growth theory*. Originally formulated decades ago (Cleland, 1971; Hager *et al*., 1971), the core still holds true: auxin apoplast acidification is induced which activates enzymes responsible for cell wall extensibility. That is to say, if apoplast acidification is not occurring, cell elongation and ultimately growth is inhibited (Kaiser & Scheuring, 2020). We hypothesize that SA impacts cell wall status (flexibility) via inhibition of TMK1 and subsequent suppression of AHA activity. By potentially acting downstream of ABP1, SA might attenuate growth independently of TIR/AFB and ABP1 auxin receptors. Intriguingly, root apoplastic pH has been described as integrator of plant signalling and several phytohormones alter the pH as consequence of their response mechanisms (Gámez-Arjona *et al*., 2022). Treatment with abscisic acid (ABA) e.g. specifically inhibits AHA activity leading to alkalinization of the apoplast and interferes with root growth (Planes *et al*., 2015). Remarkably, increase in apoplastic pH is accompanied by decrease in cytosolic pH similar to what we have determined (Fig. 1). However, low ABA concentrations seems also capable to stimulate AHA activity (Miao *et al*., 2021), suggesting a concentration-dependent and potentially cell-type specific effect.

During root cell differentiation, cytokinin activates AHAs to acidify the apoplast which then activates the α-expansin EXPA1 to loosen the cell wall (Pacifici *et al*., 2018). Notably, cytokinin has also been shown to promote root growth cessation and it seems likely that cell wall stiffening is the underlying molecular mechanism (Liu *et al*., 2022). Even more intricate, the root surface pH of Arabidopsis shows acidic and alkaline zones which are not determined by AHA activity (Serre *et al*., 2023). The root transition zone shows an alkaline pH which depends on the function of the AUX1 auxin influx carrier, the AFB1 auxin co-receptor, and the CNCG14 calcium channel. This indicates that several distinct mechanisms must be integrated to allow plant roots to rapidly change the pH of their cell walls to efficiently regulate growth.

Taken together, our data demonstrated that SA application induces apoplast alkalinization and cytosol acidification thereby attenuating Arabidopsis root growth. The SA-induced apoplast alkalinization is *TMK1* -dependent and impacts PM H^+^-ATPase function at the cell surface. Since neither nuclear TIR/AFB signalling nor the receptor ABP1 seems to be involved, this could indicate that SA-induced apoplast alkalinization functions independently of auxin signalling. However, ABP1 has been found to act redundantly with the recently identified ABP1-like (ABLs) receptors which all bind auxin and act as co-receptors with TMKs (Yu *et al*., 2023; Rodriguez *et al*., 2025). Therefore, it cannot be ruled out that loss of ABP1 is compensated by structural homologues and SA-induced alkalinization might, theoretically, involve ABLs. By potentially boarding the rapid auxin signalling pathway, SA might integrate stress responses into plant growth and developmental processes. A scenario seems feasible where specific targets of auxin-induced ultrafast ABP1-TMK1-based phosphorylation (Friml *et al*., 2022) might be blocked by SA to regulate signalling output depending on environmental cues.

## Materials & methods

### Plant Material and Growth Conditions

*Arabidopsis thaliana* (Arabidopsis hereafter) ecotype Columbia-0 (Col-0) was used as control. The lines were described previously: cpYFP (Schwarzländer et al, 2011), *npr1-1 npr3-1 npr4-3* (*npr1 npr3 npr4*) (Zhang *et al*., 2006), *pp2aa1-6* (*pp2a*) (Blakeslee *et al*., 2008), *tir1 afb2 afb3* (Parry *et al*., 2009), *tmk1-1* (Cao *et al*., 2019), *aha4-1* (Vitart *et al*., 2001) and *abp1-C1* (Gao *et al*., 2015). Seeds were surface sterilized using ethanol and stratified at 4 °C for 1–2 days in the dark. Seedlings were grown vertically at 22 °C under a 16 h light/8 h dark-cycle on half-strength Murashige and Skoog (MS) medium (Duchefa, Netherlands), including 1% (w/v) sucrose (Roth, Germany), 2.5 mM MES (Duchefa) and 1% (w/v) Phytoagar (Duchefa) at pH 5.7.

### Chemicals and Treatments

The dye HPTS was acquired from abcr (Germany) and dissolved in water. Salicylic acid was obtained from Duchefa (Netherlands) and fusicoccin (FC) from Biomol (Germany). Both were dissolved in dimethyl sulfoxide (DMSO).

### Confocal Microscopy

Live cell imaging was performed using a Zeiss LSM880, AxioObserver SP7 confocal laser-scanning microscope, equipped with either a Zeiss C-Apochromat 40×/1.2 W AutoCorr M27 water-immersion objective or a Plan-Apochromat 20x/0.8 M27 objective (INST 248/254-1).

For apoplast pH measurements based on HPTS, images were recorded using 405 nm and 488 nm excitation and 509-527 nm emission wavelengths. To determine relative cytosolic pH based on cpYFP the same excitation and emission parameters were used. Images were captured and processed using the Zeiss software ZEN 2.3 or the Fiji software (Schindelin *et al*., 2012).

### Apoplastic pH measurement using HPTS staining

For HPTS staining, seedlings were incubated for 45 minutes in liquid half-strength MS medium containing HPTS. After staining, seedlings were placed on a slide containing HPTS-supplemented medium. Fluorescence signals were detected using a Zeiss Plan-Apochromat 20×/0.80 M27 objective, capturing both the protonated (excitation 488 nm) and the deprotonated (excitation 405 nm) form of HPTS. The data was evaluated using a Fiji macro tool which detects and calculates the ratio of the 488 nm/405 nm signal (Dataset S1). Ratiometric images were generated as described previously (Barbez et al., 2017).

### Cytoplasmic pH measurement using cpYFP

For measuring cytosolic pH, the Arabidopsis marker line cpYFP was used. Fluorescence intensity ratio (488/405 nm) was calculated to determine pH changes in the cytosol using the macro described above. Ratiometric images were generated as described previously (Barbez et al., 2017).

### Phenotype Analysis

For phenotypical analysis, Arabidopsis seedlings were grown on vertically oriented agar plates containing half-strength MS medium. To analyze short-term growth, a SPIRO (Smart Plate Imaging Robot) was used (Ohlsson et al, 2024). To this end, Arabidopsis seedlings were grown for 6 days and then transferred to half-strength MS agar plates without sugar and in presence or absence (=control) of SA. Images were acquired hourly for 5 hours to determine *additional growth*. This is growth (=root length) in 5 h related to the length of the entire root after 6 days in percent. For relative root length, the tested mutant or condition was normalized to the mean root length of the corresponding control.

### Root growth rate

Seeds were surface sterilized using chlorine gas for 3 hours. For seeds culture, the RoPod5 from Guichard et al., 2024 was used. They were surface sterilized by immersion into 1% Peracetic acid, together with an exposure to ultrasound for 3 minutes, followed by 15 min incubation on a shaker. The RoPods were then washed three times with sterile water followed by an incubation of 15 minutes in sterile water. They were then sprayed with 70% Ethanol and let dry under a sterile bench, while being exposed for 1 hour to the UV lamp of this bench. 3mL of 1/2MS medium (pH5.7) supplemented with 1% sucrose and 0.8% of plant agar was added into the RoPod, and subsequently a band of 5mm of solidified medium was removed from the top of the long side of the RoPod. The seeds were positioned at the junction between the cut agar and the microscope cover-slide. The RoPod with the seeds were placed in an empty squared dish, stratified at 4°C in the dark for 2 to 3 days, and positioned in an angle of 45° in a culture cabinet set at 22°C and 16h illumination, for 5 to 6 days. The RoPods were installed on a vertical microscope (Zeiss AxioVert 200M, ITK MMS-100 linear stage and Hydra controller, Hamatsu OrcaC11440-36U camera, Zeiss 10 × PlanNeoFluar objective). LED bulb (LED power 150W, Chip SMD2835, 215 pcs Warm, 40pcs Red, 5pcs blue) was used for bright field illumination. The root images were acquired every 2.8 minutes. A base line of 3 hours without any treatment was performed. Subsequently, 1/2 MS medium supplemented with 1% sucrose and a mock solution (absolute ethanol) or 70µM SA (for a final concentration of 50µM SA after equilibration in the RoPod) was injected. Root growth was recorded over the following 5 hours. The obtained images were analyzed using ImageJ (Schindelin *et al*., 2012). They were first converted to a 8-bit image, and when needed stitched and registered. Afterward, the root tips were tracked using TrackMate (Tinevez *et al*., 2017). The generated measurements were analyzed with R.

### PM H^+^-ATPase activity assay

To measure PM H^+^-ATPase activity, seedlings were grown on half-strength MS medium for 4 days and transferred to liquid ½ MS medium for two weeks. The seedlings were treated with 50 µM SA for 5 hours and with 1 µM FC for 1 hour, or corresponding control solvents (DMSO and EtOH). Root samples were isolated and frozen in liquid nitrogen. Proteins were isolated in homogenization buffer (50 mM Tris-HCl, pH 7.5, 150 mM NaCl; 10% glycerol, 2 mM EDTA, 5 mM DTT, 1 % protease inhibitor cocktail, 1 % IGEPAL, 1 mM NaF, 1 mM PMSF). Protein samples were clarified by centrifugation, 13 000 x g for 20 min at 4 degrees. Microsomal fractions were collected from supernatants by ultracentrifugation at 100 000 x g for 1 hour at 4 degrees. The microsomes were resuspended in homogenization buffer to a protein concentration of 0.45 µg/µl. The ATP hydrolytic activity of vanadate-sensitive PM H^+^-ATPase were measured by quantifying phosphate-malachite complexes as a proxy for the released inorganic phosphate as described in (Oecking *et al*., 1997).

### Immunoblotting

Protein samples were separated in 10 % bisacrylamide gels at 150 V for approximately 2 hours and transferred into activated PVDF membranes at 100 V for 60 min. Immunoblotting was performed in blocking solution (5% milk in TBS with 1% [v/v] Tween-20) with the following antibodies α-H^+^-ATPase (1:3000, Agrisera AS07260), α-pAHA2 (1:5000, Agrisera AS22 4789), α-14-3-3 (1:2000, Agrisera AS122119), α-cAPX (Agrisera AS06180), and the secondary anti-rabbit IgG (whole molecule)–HRP (A0545, Sigma, dilution 1:10,000). Western blots were developed with SuperSignal™ West Pico PLUS Chemiluminescent Substrate (Thermo Scientific).

### Statistical analysis

All quantitative data was analyzed using the GraphPad Prism 9 software. The precise statistical method used are given in the respective figure legends.

## Supplemental data

Figure S1: *tmk1-1* but not *aha4-1* is partially resistant to SA-mediated root growth inhibition. Figure S2: *abp1-c1* is not resistant to SA-mediated root growth inhibition.

Dataset S1

## Acknowledgements

For providing published material we are grateful to Jiri Friml

We thank Matthias Hahn for critical reading of the manuscript. This work was supported by the Deutsche Forschungsgemeinschaft (DFG) grants CRC1101-A09 and TRR356-B01 to J.G., a DFG Heisenberg Professorship (GR4559/4-1) and the German Excellence Strategy (CEPLAS-EXC-2048/1-project ID 390686111) to G.G., and grants from the *BioComp* research initiative (Rhineland-Palatinate, Germany) and the DFG (SCHE 1836/4-2 and SCHE 1836/5-1) to D.S.

## Contributions

J.M. performed most experiments. K.X., M.G., S.K., R.T. and D.S. carried out and analysed experiments. J.M., G.G., J.G. and DS participated in developing the experimental setup. J.M. and D.S. designed the figures and performed statistical analysis. D.S. conceived the study and J.M. and D.S. wrote the manuscript. All authors saw and commented on the manuscript.

