## Supplementary Data for "Salicylic acid-induced alkalinization of the apoplast requires *TRANSMEMBRANE KINASE 1* and results in growth attenuation"

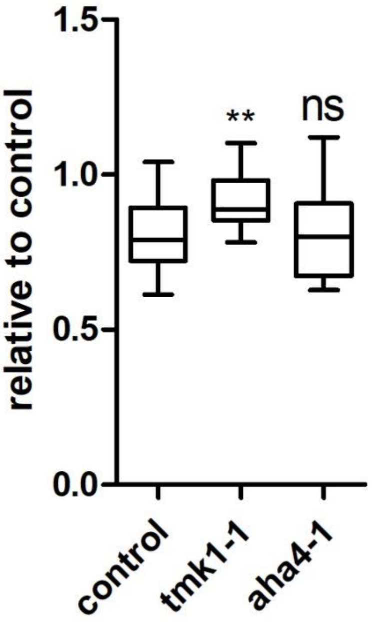


**Figure S1. *tmk1-1* but not *aha4-1* is partially resistant to SA-mediated root growth inhibition.** Six-day-old seedlings of Col-0, *tmk1-1*, and *aha4-1* were grown on ½ MS- plates and then either transferred to ½MS- plates or ½MS- plates containing 50 µM SA. Root growth was measured after 5 hours, expressed relative to the total root length and then normalized to its given (mock-) control (n Col-0 = 48, n Col-0 SA = 49, n *tmk1-1* = 34, n *tmk1-1* SA = 37, n *aha4-1* = 36, n *aha4-1* SA = 32). This figure is based on the data from Fig. 3C. Data is presented in whisker plots with statistical analysis performed using Student's t-test. **p≤0.01.

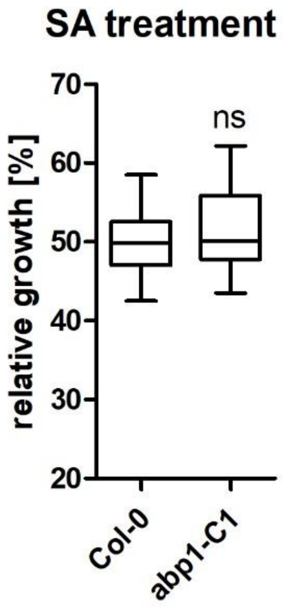


**Figure S2. *abp1-c1* is not resistant to SA-mediated root growth inhibition.** Root growth analysis of Col-0 and *abp1-c1* seedlings grown for 7 days on ½ MS+ plates containing 50 μM SA. Relative growth was calculated as the ratio of root length on SA-supplemented media to the average root length of seedlings grown on control media (n Col-0 = 22, n *abp1-c1* = 18). This figure is based on the data from Fig. 4C. Data is presented in whisker plots with statistical analysis performed using Student's t-test. ***$p \leq 0.001$.

**Dataset S1.**

Upon opening the macro, the variable “setThreshold” can be adjusted to match the signal intensity.

This macro quantifies the mean gray value of both channels of a ratiometric dye. It goes through each image of the chosen folder, splits the channels into separate images, renames them according to their respective wavelengths, applies the defined threshold, and measures the mean gray value of the threshold-outlined signal.

The macro requires the FeatureJ plugin in Fiji. It is recommended to use the Fiji Life-Line version 1.52u (17. March 2020) for full compatibility.

The output consists of two values per image, corresponding to the mean gray values of the two measured wavelengths.

```
// Ask for Bio-Formats
Dialog.create("Normalization of Marker");
Dialog.addCheckbox("Use Bio-Formats Importer ", true);
Dialog.show();
Bio=Dialog.getCheckbox();

input = getDirectory("location where images are stored");
list = getFileList(input);

setBatchMode(false);

// Loop to sequentially open images
for (i=0; i<list.length; i=i+1)
{
   full = input + list[i];
   print(full);

   if (Bio==true){
      run("Bio-Formats Importer", "open=&full autoscale color_mode=Composite
view=Hyperstack stack_order=XYZCT");
   }
   else {
      open(full);
   }
}

// Get the name without the extension
filename=getTitle();
ShortFileName=substring(filename, 0, lastIndexOf(filename,"."));

// Split & rename
run("Split Channels");
selectWindow("C1-" + filename);
rename("488");
selectWindow("C2-" + filename);
rename("405");
selectWindow("488");
run("Duplicate...", "title=RFP-1");
```

```
run("Mean...", "radius=2");
setThreshold(15, 250);
run("Create Selection");
run("Add to Manager");
close();
selectWindow("488");
roiManager("Select", 0);
run("Measure");
run("Select None");
selectWindow("405");
roiManager("Select", 0);
run("Measure");
run("Select None");
roiManager("Delete");
selectWindow("405");
run("Close All");
```
